# Modeling population size independent tissue epigenomes by ChIL-seq with single-thin sections

**DOI:** 10.1101/2020.12.18.423434

**Authors:** Kazumitsu Maehara, Kosuke Tomimatsu, Akihito Harada, Kaori Tanaka, Shoko Sato, Seiji Okada, Tetsuya Handa, Hitoshi Kurumizaka, Hiroshi Kimura, Yasuyuki Ohkawa

**Author notes:** Corresponding author: Yasuyuki Ohkawa, **Email:**.

## Abstract

Recent advances in omics studies have enabled analysis at the single-cell level; however, methods for analyzing the whole cell of large organs and tissues remain challenging. Here, we developed a method named tsChIL to understand the diverse cellular dynamics at the tissue level using high-depth epigenomic data. tsChIL allowed the analysis of a single tissue section and could reproducibly acquire epigenomic profiles from several types of tissues, based on the distribution of target epigenomic states, tissue morphology, and number of cells. The proposed method enabled the independent evaluation of changes in cell populations and gene activation of cells in regenerating skeletal muscle tissues, using a statistical model of RNA polymerase II distribution on gene loci. Thus, the integrative analysis by tsChIL can elucidate *in vivo* cell-type dynamics of tissues.

## Introduction

Tissues are terminally differentiated cells formed from stem cells, followed by cell-type conversion and functional arrangement of cell types to the specified spatial localization. Presently, the composition of the cells playing different functions and the mechanism by which each type is formed have been elucidated. This allowed us to understand the biological function of each tissue and the pathogenesis and developmental failure of diseases. Tissue composition can be determined using known cell-type markers. Immunostaining for cell surface antigens and other cell-type markers enables visual examination that determines the number and localization of cells and tissue morphology. Determining the cell types and the size of the population (i.e. number of cells) in tissues can be done using scRNA-seq, which is based on the transcriptomic differences of individual cells^1^. Unsupervised clustering^2^ of the gene-expression profiles allowed the identification of the cell types, including those previously unknown.

Epigenomic analysis is widely performed at the tissue level, such as in the Encyclopedia of DNA Elements (ENCODE), the National Institutes of Health Roadmap Epigenomics Project, and International Human Epigenome Consortium (IHEC) projects that utilize ChIP-seq for bulk-level tissues. The comprehensive identification of functional elements in genomes, such as promoters, enhancers, and the binding sites of transcription factors and their regulatory relationships that characterize the tissues has been achieved^3–5^. However, in epigenomic analysis at tissue-level, imbalance sampling cannot be avoided because tissues are mixtures of diverse cell types. Particularly, when the number of target cells (e.g. stem cells) is limited, they are overwhelmed by the information on the other majority cells. Furthermore, in ChIP-seq, the genome coverage per cell is limited^6^, i.e. information on minority cells in the bulk tissue is lost. Therefore, it is necessary to collect a large amount of target cells that meet the requirements of ChIP-seq. As such, after defining the target cell types and markers, the sectioning of narrower area, dissection, or cell-sorting is utilized. Recently, new methods for analyzing a small number of cells with higher genome coverage at the single-cell level, including our ChIL and others, have been developed^7–14^. In addition, isolating cells causes potential effects to cells owing to the physical separation of the tissues. Several tissue analysis methods that do not involve enzymatic digestion, have also been proposed^15–20^. However, obtaining epigenomic information from a limited number of cells using ChIP-seq based technology remains a challenge. To understand the formation of all cell types, the use of whole-tissue analysis using single-cell technology is ideal; however, it is very costly.

Hence, several transcriptomic analysis approaches that combine the advantages of bulk RNA-seq and scRNA-seq, which can analyze and identify numerous cells at once, have been proposed. For example, the changes in the gene expression of cell types in bulk RNA-seq profiles during embryogenesis have been interpreted using single cell RNA-seq collected separately^21^. The estimation of the cell-type composition of bulk tissue RNA-seq based on single cell RNA-seq has also been reported^22^. Because data from different platforms complement each other, a data integration method has also been proposed, particularly the embedding of single cell RNA-seq into seqFISH+^23^ data to virtually reconstruct whole gene expression data using spatial information^24,25^. In addition, a computational approach for epigenomic analysis to decompose DNA methylation states into cell types has been suggested^26^. However, to date, a universal solution for the cell-type decomposition of tissue epigenomes has not yet been established.

Here, we propose a framework that integrates ChIL into the tissue-slice analysis and uses single, very small, and thin tissue sections for ChIL-Seq (herein, tsChIL). Our obtained bulk tissue epigenome data showed dynamic changes in both the number and cell type, and computational modeling was thus required. We first optimized ChIL for highly sensitive genome-wide analysis using a single thin section, as well as tissue visualization using immunostaining. The ChIL is proposed to enable epigenomic analysis at the single-cell level, and the acquired thin-section tsChIL data are expected to be the sum of the high-depth single-cell epigenomes. Using three different types of tissues, we confirmed the adequate sensitivity, specificity, and reproducibility of tsChIL in identifying enhancers, transcription factors, and transcriptionally activated genes in whole tissues. Thus, we built a statistical model that evaluates the changes in the distribution of RNA Polymerase II at the gene loci and provides a robust, cell-type resolution transcriptional regulatory analysis for large changes in population size.

## Results

### Thin-section ChIL-seq enabled spatial epigenomics with single tissue section

Various cell types exist in tissues, each of which exhibits a unique localization pattern. The transcriptomic and epigenomic pattern of these cells may be affected by the enzymatic isolation process. Therefore, we focused on the use of tissue sections that are free from enzymatic treatment biases for epigenomic analysis and developed an experimental procedure using a single, very small, and thin tissue sections. We then optimized the ChIL for tissue **(Fig. 1A)**, based on our previously reported sc-epigenomic analysis tools^7,8^.

**Figure 1:**
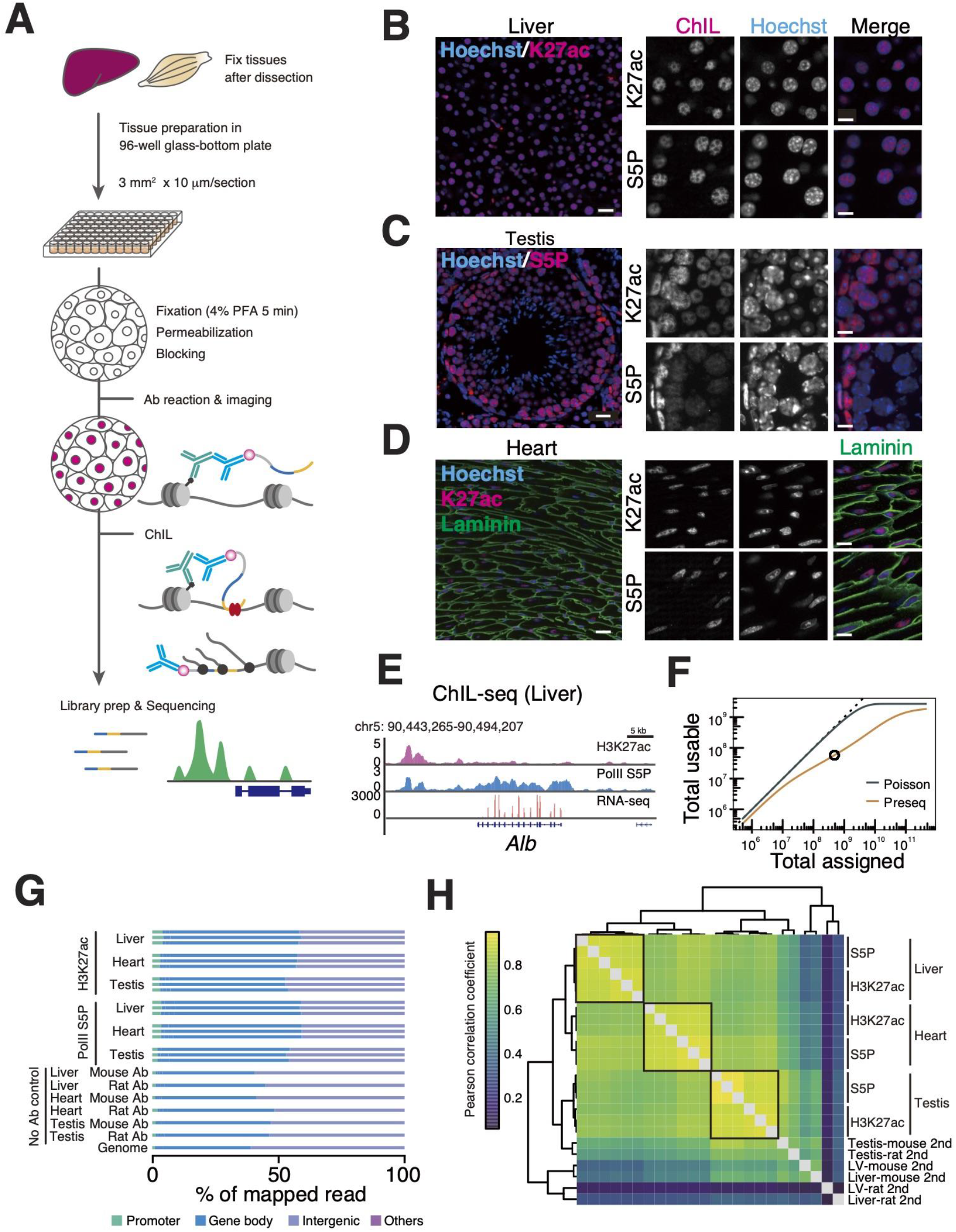
Epigenomic profiling using a single tissue section. **(A)** Schematic diagram of the tsChIL protocol. **(B-D)** Immunofluorescent images of mouse liver **(B)**, testis **(C)** and heart **(D)**. H3K27ac, RNAPII-S5P, and laminin were stained with specific primary antibodies and visualized using fluorescent labeled anti-mouse ChIL-probe (red: H3K27ac and PolIIS5P) and anti-rabbit IgG (green: laminin). DNA was counterstained with Hoechst 33342. Scale Bar: 20 μm (left images), 10 μm (right images). **(E)** Genome browser images of ChIL-seq for H3K27ac and RNAPII-S5P and bulk tissue RNA-seq data at the *Alb* locus in liver tissues. **(F)** Library complexity of ChIL data. Poisson represents an ideal case of the uniform probability of obtaining reads from the mouse genome, whereas preseq refers to the future/past predictions of a species discovery curve of sequenced reads using Preseq^28^. Black circle indicates the read number we sequenced for this prediction. **(G)** Breakdown of mapped reads at the annotated genomic regions. Gene body: 3’-UTR, exon, intron, 5’-UTR; Others: ncRNA, miRNA, snoRNA, and pseudogenes. The proportions of the annotated region on the mouse genome are shown as “Genome” at the bottom lane. **(H)** Genome-wide correlation at 10 kbp bins. Hierarchical clustering of Pearson’s correlation coefficient of log-transformed tsChIL-seq counts is shown.

Since the reports on analysis using microtissue sections are limited, and all of them require multiple tissue slices to obtain the required cell number^15–20^ (**Table S1**). We therefore focused on preparing frozen, unfixed tissue sections to equalize the fixation conditions. On plates, unfixed tissue thin sections are fixed with paraformaldehyde then permeabilized, followed by blocking. Immunostaining is then performed by reacting with primary antibodies against the target molecules on chromatin. Then, a fluorescent-labeled ChIL-probe attached with secondary antibodies was used to obtain the tissue localization of the target by imaging at the subcellular level. Subsequently, Tn5 transposase inserts an artificial sequence containing a T7 promoter into the genomic region near the target. In vitro transcription of the genome sequence near the target protein, starting from the T7 promoter, was performed, and the reverse-transcribed DNA was sequenced using a next-generation sequencer. Compared with conventional epigenomic analysis methods for FFPE and fresh frozen tissue slices, this method enabled uniform fixation conditions for the analysis of micro-thin slices. Therefore, using the highly efficient ChIL method, we attempted to analyze tissues with an input size of 3 mm × 3 mm × 10 μm. Thus, we designed tsChIL as a high-precision method for analyzing the epigenetic information of a group of cells on a tissue section of the target, following the spatial distribution of the specific epigenetic status.

To evaluate the designed tsChIL experimental procedure, the levels of the enhancer marker of histone modification H3K27ac and the recruitment of RNA Polymerase II (RNAPII), an indicator of transcription, were detected in three different tissues: liver, heart (left ventricle), and testis. Most of the cells were hepatocytes, comprising 70–80% of the liver. The H3K27ac signal visualized by the ChIL-probe was uniformly distributed across cells on the sections. Subcellularly, the co-localization of H3K27ac and RNAPII in euchromatin regions (Hoechst-negative) was observed (**Fig. 1B**). In the testis, which consists of cells at multiple differentiation stages, the RNAPII signal was strongly distributed and localized in cells with high transcriptional activity, especially near the outer periphery of the seminiferous tubule^27^ (**Fig. 1C**), a region where cells in the early stages of sperm differentiation are located **(Fig. 1C)**. Meanwhile, the heart was co-stained using laminin and the ChIL probe to distinguish the cell boundary regions and visualize the basement membrane (**Fig. 1D**). S5P signal showed a localization to the low Hoechst-dense region of the cell nucleus in which transcription may active, suggesting that immunostaining with ChIL probe was a valid histological staining method at the subcellular level (**Fig. 1B-D, Fig. S1**).

To validate the feasibility of tsChIL for sensitive and accurate epigenomic analysis, we performed tsChIL-seq using a single thin section containing 1,000-10,000 cells (**Table 1**), which was generally assumed as a low number of cells in culture^7,8^. The number of cells used was less than that of conventional epigenomic methods used especially for tissue analysis (**Table S1**). Furthermore, the genome-wide analysis was performed by ChIL reaction on single sections of the sections that showed in **Figure 1B-D**. In the representative visualized epigenomic data in liver (**Fig. 1E)**, the accumulation of H3K27ac and RNAPII at the *Alb* locus, a hepatocyte marker, was observed. The former showed an activated upstream enhancer region, whereas the latter was highly transcriptional activity at the locus. The transcription of *Alb* was also confirmed using RNA-seq with different serial slices. These results indicate that tsChIL enables the simultaneous acquisition of both the tissue distribution of the epigenomic status and the genome-wide epigenomic data using a single tissue section containing a small number of cells (10^3^ to 10^4^ cells in 10 mm^2^ area).

**Table 1:**
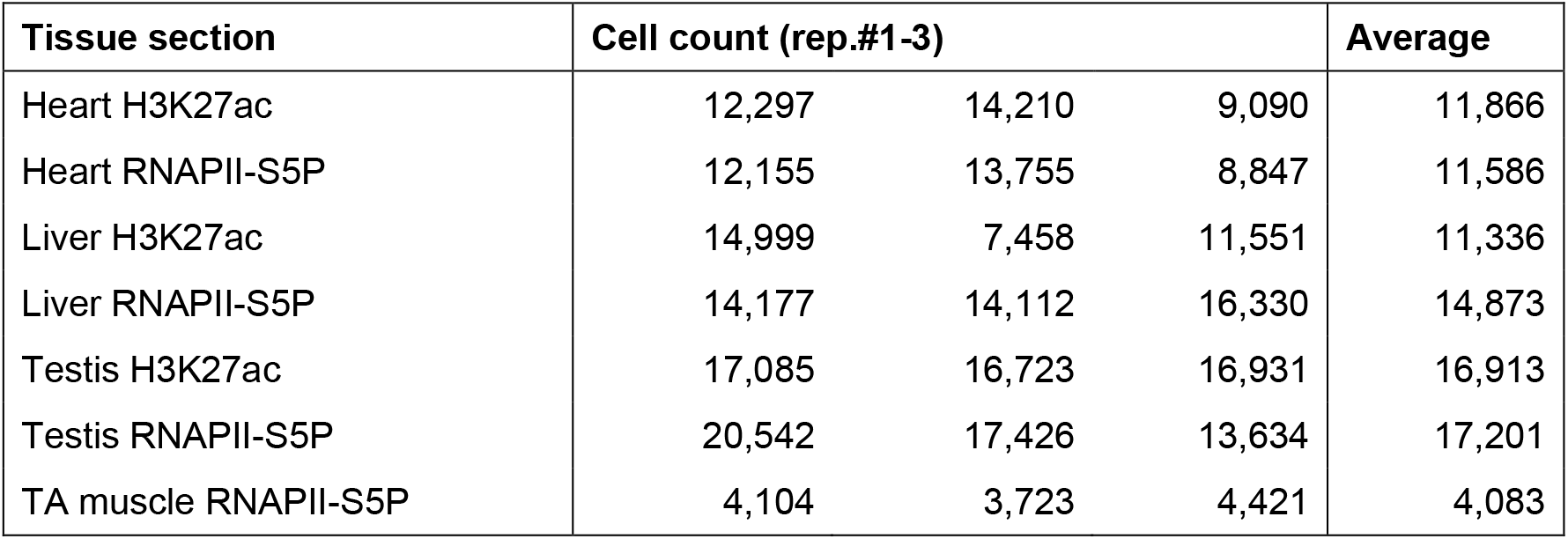
Cell numbers in the tissue sections used in this study.

Next, to evaluate the genome-wide distribution of the signals obtained using the tsChIL procedure proposed above, we examined the specificity of the signal localization among different tissues and antibodies and the reproducibility of signal localization of the same tissue and antibody. First, to estimate the appropriate number of reads for ChIL-seq with tissues, we obtained 480 M reads from RNAPII ChIL-seq in muscle tissue and evaluated the library complexity^28^ (i.e. the prediction curve of usable reads). As seen in **Figure 1F**, the number of total usable reads was starting to move away from the black line at approximately 10^7^, indicating a decreasing percentage of usable reads. Therefore, we determined that approximately 10^7^ reads is a good cost-balanced number of the required reads in the case wherein the number of cells per section is < 10^4^. To obtain a ChIL signal with sufficiently high signal-to-noise ratio, we acquired an average of approximately 14 M reads (**Table S2**), which is comparable to the number of reads in the ENCODE tissue ChIP-seq (10 M–20 M)^3^.

With this number of reads, the tsChIL-seq data from the liver, heart, and testis were obtained, and the genome-wide localization of each data set is shown in **Figure 1G**. In all tissues and H3K27ac and RNAPII S5P antibodies, signals were concentrated around the coding regions (promoters and gene body) compared with the no-antibody (herein, No Ab; without primary antibody) controls (53%-59% and 41%-48%, respectively). The results showed that the genomic sequences were selectively extracted from the transcriptionally activated regions of the genome. In **Figure 1H**, we describe the correlation matrix of the signal levels on the whole genome to confirm the high reproducibility of the replicates. The dendrogram shows the hierarchical structure of the highest correlation among the replicates (Liver-H3K2ac: 0.90, Liver-PolII: 0.90, Heart-H3K27ac: 0.87, Heart-PolII: 0.92, Testis-H3K27ac: 0.91, Testis-PolII: 0.94 in average of triplicates), and the correlation within the same tissue (e.g., Liver-PolII vs. Liver-K27ac: 0.87; Heart-H3K27ac vs. Heart-PolII: 0.88; and Testis-H3K27ac vs. Testis-PolII: 0.88; the list of all correlation coefficients are summarized in **Table S3**). These results suggest that tsChIL-seq can capture the epigenomic differences between different tissues and is technically reproducible.

### Identification of regulatory factors in the formation of tissue-specific enhancers

We next assessed the ability of tsChIL for low-input epigenomic analysis of tissues. First, we performed tsChIL using thinly sectioned tissues from the liver, heart, and testis, and the identified enhancers were compared by matching references^3^ (**Fig. 2A**). According to the odds ratio (i.e., specificity, the detailed definition is described in Method), each H3K27ac ChIL-seq signal preferentially captured the corresponding tissues-specific enhancer (Liver: 33.5, Heart: 27.1, Testis: 4.1; The other odd ratios are listed in **Table S4**). Therefore, we successfully detected tissue-specific enhancers using tsChIL with lower input compared to the previous reports that utilized 500 μg chromatin equivalent to 10^7^-10^8^ cells.

**Figure 2:**
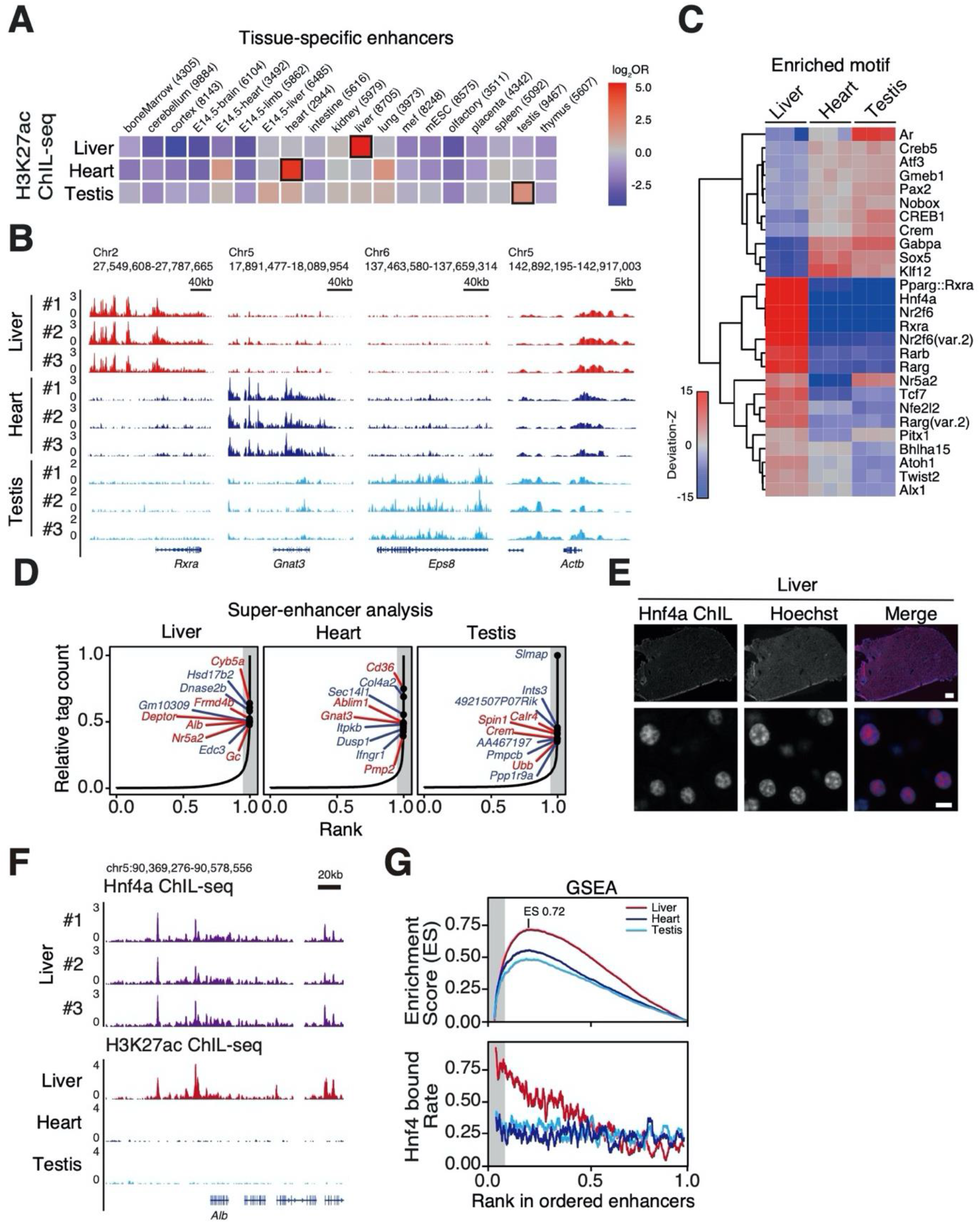
Upstream factor identification through enhancer analysis using tsChIL-seq. **(A)** Tissue specificity of identified enhancers by tsChIL-H3K27ac. The odds ratios of hits in the reference tissue-specific enhancer list identified by bulk-tissue ChIP-seq data^3^ are shown. Odds is defined in Methods. The cells enclosed by black squares indicate the maximal odds ratios (i.e., maximal specificity) for each row. **(B)** The IGV tracks of tsChIL-H3K27ac at identified tissue-specific enhancers of *Rxra, Gnat3, Eps8*, and a house-keeping gene of *Actb* loci are shown with the replicates. **(C)** Specific motif enrichment analysis was conducted using chromVAR^66^. Hierarchical clustering of deviation-Z scores of three replicates of each tissue is shown. **(D)** Super-enhancer identification. Tissue-specific enhancers are identified, so that they are listed more than twice (twice: blue, all: red) in the top 5% in all enhancer candidates and are not in the SEs of other tissues. Grey shades indicate the top 5% of tag count in enhancer candidates. **(E)** Immunofluorescent images of mouse liver sections. Tissues were stained with anti-Hnf4α antibody and visualized by a fluorescent-labeled anti-mouse ChIL probe. DNA was counterstained with Hoechst 33342. Scale bar: 200 μm (top), 10 μm (bottom). **(F)** Hnf4α binds to the SE at *Alb* gene loci. **(G)** Gene set enrichment analysis of Hnf4α-bound genes (top), and their rate of Hnf4α-bound genes in sliding windows of 100 genes (bottom).

Next, we examined the enrichment of the H3K27ac signal on representative tissue-specific enhancers, including the liver, heart, and testis. We focused on the enhancer region of *Rxra* genes^29^ specifically expressed in liver tissues, *Gnat3* cardiac muscle-specific gene retinoic acid receptor, and *Eps8* expressed in the blood–testis barrier (BTB)^30^. H3K27ac signal enrichments on each tissue-specific enhancer were observed on the IGV screen shot (**Fig. 2B).** In contrast, *all Actb*-expressing tissues showed the ubiquitous enrichment of H3K27ac.

We further evaluated the enrichment of the regulatory sequence in extracted enhancers using tsChIL based on the enrichment of the TF-binding motif (only the top scoring motifs are shown in **Fig. 2C**; all others are in **Table S5**). The enrichment of known liver-specific TF-binding motifs, Rxra, Hnf4a, Nr2f6, and others were observed in the H3K27ac tsChIL-seq data obtained from the liver. This data is consistent with the liver-specific regulatory sequences registered as open chromatin regions detected using ATAC-seq with mouse liver tissues in the database^31^. Meanwhile, the H3K27ac signal obtained from the heart showed relatively higher enrichment at Klf12 than others; Sox5 and androgen receptor (AR) binding motifs were enriched in the testis-H3K27ac signal, which was consistent with previous studies reporting that AR binds to the androgen responsible element (ARE) on regulatory sequences with histone acetyltransferase to regulate gene expression^32,33^. These data support that H3K27ac tsChIL can identify *cis*-regulatory elements following the extraction of tissue-specific enhancers.

Because the enrichment of the tsChIL signal should reflect the quantitative H3K27ac levels as demonstrated by the identification of super enhancers (SEs) using ChIP-seq, we next quantitatively determined the H3K27ac level based on the read counts. Then, SE formation upon TF binding on the extracted cis-regulatory elements was evaluated. First, we listed the highly enriched regions of the H3K27ac tsChIL signal as SE^34,35^ from each liver, heart, and testis data set. The labeled genes in **Figure 2D** are representative protein-coding genes near the identified top ranked SEs, which have the highest read counts in peaks (see **Fig. S2** for all replicates). In the liver, known hepatocyte marker genes, *Alb*, and albumin family, *Gc* are also detected in motif-enrichment analysis performed in **Figure 2C**. In addition, the core transcription factor Hnf4α^36^, which activates the genes by itself, was included in the top rank (1.6 to 3.5%). Furthermore, the SEs featuring each tissue were identified. In the heart (left ventricle), *Ablim1* expressed in the left ventricle and involved in left–right axis formation^37^, was detected, whereas in the testis, SEs were identified in the vicinity of *Crem*, which is involved in spermatogenesis^38^.

Finally, to validate the function of the SEs identified in the liver using this method, we performed tsChIL targeting Hnf4α, which showed a high specificity score (deviation-Z) in liver SEs. Hnf4α is known to be an important nuclear receptor during hepatocyte differentiation^39^, and has been shown to contribute to SE formation as a core transcription factor, along with RXRα^29^. Immunostaining with the ChIL Probe showed that the HNF4α was distributed throughout most cells in the liver tissue and detected in the open chromatin region of the nucleus in each cell (**Fig. 2E**). A pronounced accumulation of Hnf4α signals in the SEs in the region was observed (**Fig. 2F**, see **Fig. S3** for the motif enrichment analysis on Hnf4α peaks). We next evaluated the selective binding of Hnf4α to the genes in the liver SEs (**Fig. 2G**; **Fig. S4** for the replicates). Using the gene sets of SEs and TEs neighboring genes obtained in **Figure 2D**, gene sets enrichment analysis (GSEA)^40^ demonstrated that the hits of the ChIL-Hnf4α peaks against liver enhancers scored as high as 0.72 in the enrichment score (**Fig. 2G**, top). Particularly, Hnf4α was bound to 76.4–78.8% of the SEs (**Fig. 2G** bottom). In contrast, in the negative controls of the heart- and testis-specific SEs, the number of SEs bound by Hnf4α was approximately 0.5 in the enrichment score and the percentage of Hnf4α bound to the heart- and testis-specific SEs was at a random chance level (24.4–34.2%).

In summary, the data from tsChIL-H3K27ac demonstrated that the regulatory candidate transcription factor Hnf4α obtained from the *cis*-element refinement selectively binds to the liver-specific SE region of the *Hnf4a* locus. Hnf4α could be validated to provide positive feedback that binds to the SE region of its own *Hnf4a* locus. Our data indicated that tsChIL is useful for the regulatory analysis of enhancers, including transcription factors and SEs, using low number of cells.

### tsChIL-RNAPII peaks detected the majority of active genes in tissue

Transcriptome information is obtained by evaluating the binding position of RNAPII using epigenomic analysis. Here, we detected the active genes based on the binding of RNAPII on the genome using tsChIL. In **Figure 3A**, we plotted the cumulative number of consumed reads of the detected genes in RNA-seq and RNAPII tsChIL in the order of their read counts. Due to the wide dynamic range of RNA-seq data, high copy-number mitochondrial-derived RNAs (e.g., mitochondrial ribosomal RNAs) and highly expressed genes that characterize each tissue (*Alb* in liver, *Myh6* in the heart, *Prm1* in testes), consumed 80% reads on a small number of highly expressed genes (whose expression can be confirmed; Liver 5%, Heart 1%, Testis 11%). The identification of weakly expressed genes and rare populations in bulk tissue RNA-seq is generally hard to obtain because the top 10% genes spends 80% of its reads in even at the single-cell level^41,42^.

**Figure 3:**
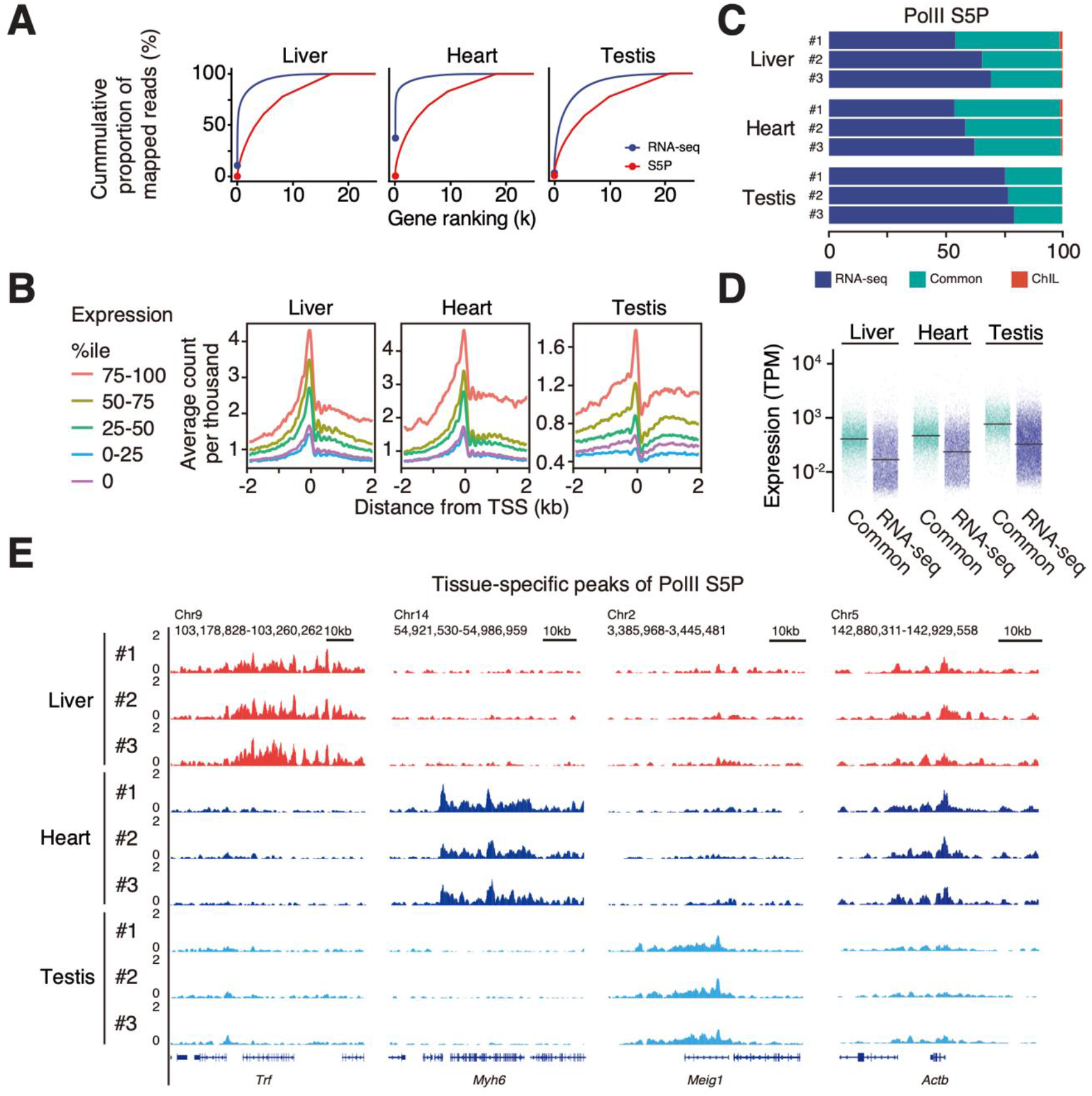
ChIL-RNAPII detect active genes in tissue. **(A)** Dynamic ranges of bulk-tissue RNA-seq and tsChIL RNAPII. The cumulative proportion in total mapped reads at genes (red: tsChIL, blue: RNA-seq) were compared. Genes are ordered by the read counts on the exons for RNA-seq and on +/-750 bp from TSS for tsChIL, respectively. **(B)** Signal intensities of tsChIL correlated with the expression levels of genes. The lines indicate the average CPM of each expression group at TSS. The expression groups were assigned with respect to the expression levels (TPM) of genes. **(C)** Coverage of expressed genes by tsChIL-RNAPII peaks. The stacked bar chart shows the proportions of detected genes in the RNA-seq only (RNA-seq: blue), tsChIL-Pol2 only (ChIL: red) and both (Common: green). **(D)** Higher expression levels at tsChIL-Pol2 peaks. The expression levels of all expressed genes (TPM > 0) are shown. **(E)** The tissue-specific genes identified by tsChIL-RNAPII. The IGV tracks of all replicates of tsChIL-RNAPII are shown at each specific gene (*Trf, Myh6*, and *Meig* for the liver, heart, and testis, respectively). *Actb* is also shown as the ubiquitously expressed gene in the three tissues.

In contrast, ChIL-RNAPII did not exhibit an exponential increase in the number of consumed reads required to detect gene expression from RNA-seq. It also efficiently detected more genes as the number of reads increased. The dynamic range of RNA-seq depends on the product of the cell number and the concentration of RNA in each gene, whereas that of the RNAPII signals, in essence, depends on the product of the presence or absence of gene expression (0, 1, or 2) and the cell number. The results are consistent with the fact that highly and ubiquitously expressed genes occupy a high number of reads in the RNA-seq data. The result suggested that fewer reads are required for gene expression profiling using tsChIL RNAP2 than RNA-seq.

Thus, the genes were divided into five groups based on their expression levels from RNA-seq, and the correlation of each tsChIL RNAPII signal with their expression levels was examined (**Fig. 3B**). In the high-expression group in all tissues, the intensity of the RNAPII signal in the TSS was highly correlated with its expression level. In the 75^th^–100^th^ percentile group, a high accumulation of RNAPII in the gene body region was also detected, suggesting a movement of RNAPII to the locus upon transcriptional activation. Here, we showed that tsChIL-RNAPII demonstrated a preference for capturing highly expressed genes in tissues. Subsequently, we assessed the overlap between RNA-seq-confirmed genes (TPM > 0) and tsChIL-RNAPII peaks. tsChIL peaks captured approximately 30% (Testis slightly lower, approximately 20%) of the active genes (TPM > 0), whereas false positives were almost absent (**Fig. 3C)**. In addition, tsChIL peaks stably detected approximately 40–50% of the genes expressed in RNA-seq, independent of the TPM threshold for defining the expressed genes in RNA-seq (**Fig. S5**). These results suggest that the peak region is likely to capture genes with high expression because the region with high signal counts was judged to be the peak region^43^. In all tissues, the expression levels of the genes in Common were higher than those in RNA-seq group as expected (**Fig. 3D**).

**Figure 3E** shows an IGV screenshot of the tsChIL RNAPII. The accumulation was detected at the *Trf* (transferrin) locus in the liver, *Myh6* (cardiac myosin) in the heart, and *Meig1* (a meiosis-expressed gene) in the testes. These are considered representatives of genes specifically expressed in each tissue. At the *Actb* locus, a house-keeping gene, the RNAPII signal was accumulated in all tissues, indicating active transcription. In these highly transcriptionally active genes, a wide distribution of RNAPII signals was detected on the gene body, suggesting that the RNAPII binding distribution patterns would enable an in-depth profiling of the transcriptional programs in tissues.

### Modeling RNAPII traveling reveals transcriptional dynamics in the rapid change of cell population in skeletal muscle regeneration

We demonstrated that enhancers and transcriptional activity states can be detected with high sensitivity, specificity, and reproducibility at the whole-tissue level by the optimized ChIL for tissues. Then, tsChIL-RNAPII data in Figure 3 suggested that, in addition to amount of the signal at the gene loci, evaluation of the distribution or its elongation across the entire locus would improve the analysis of the transcriptional activation in various cells in tissue. We thus conceived a concept the statistical modeling of tsChIL-RNAPII data for the epigenomic analysis of heterogeneous tissues.

We used skeletal muscle regeneration as a model case, wherein numerous cell types dynamically change their composition, particularly that of the mouse tibialis anterior (TA) muscle after cardiotoxin (CTX)-induced injury. During regeneration, migrating immune cells are dominate the tissue 2 to 3 days after muscle injury^44^. During this time, the activation of satellite cells, which are responsible for skeletal muscle regeneration, leads to the regenerated muscle fibers observed on day 14. We thus established a model to analyze the gene expression dynamics in each cell type from day 0 (pre-injured period) and until day 14. tsChIL obtained data from five biological replicates using the tissue sections of TA muscles at five time points on days 0, 3, 5, 7, and 14 after the CTX-induced muscle injury. As shown in **Figure 4A** (**Fig. S6A** for the entire time-course), the basal lamina separating the muscle fibers observed on day 0 was destroyed post-injury. The destruction of the cells on the third day can be seen in the image of laminin co-stained with the ChIL probe. Furthermore, the fluorescence image of the ChIL probe suggests the presence of multiple cell types, such as the activated muscle satellite cells, muscle progenitor cells that have started to differentiate, and migrating immune cells associated with the inflammatory response. On day 14, the structure of the muscle fibers possessing central nuclei were observed, thus indicating regenerated muscles.

**Figure 4:**
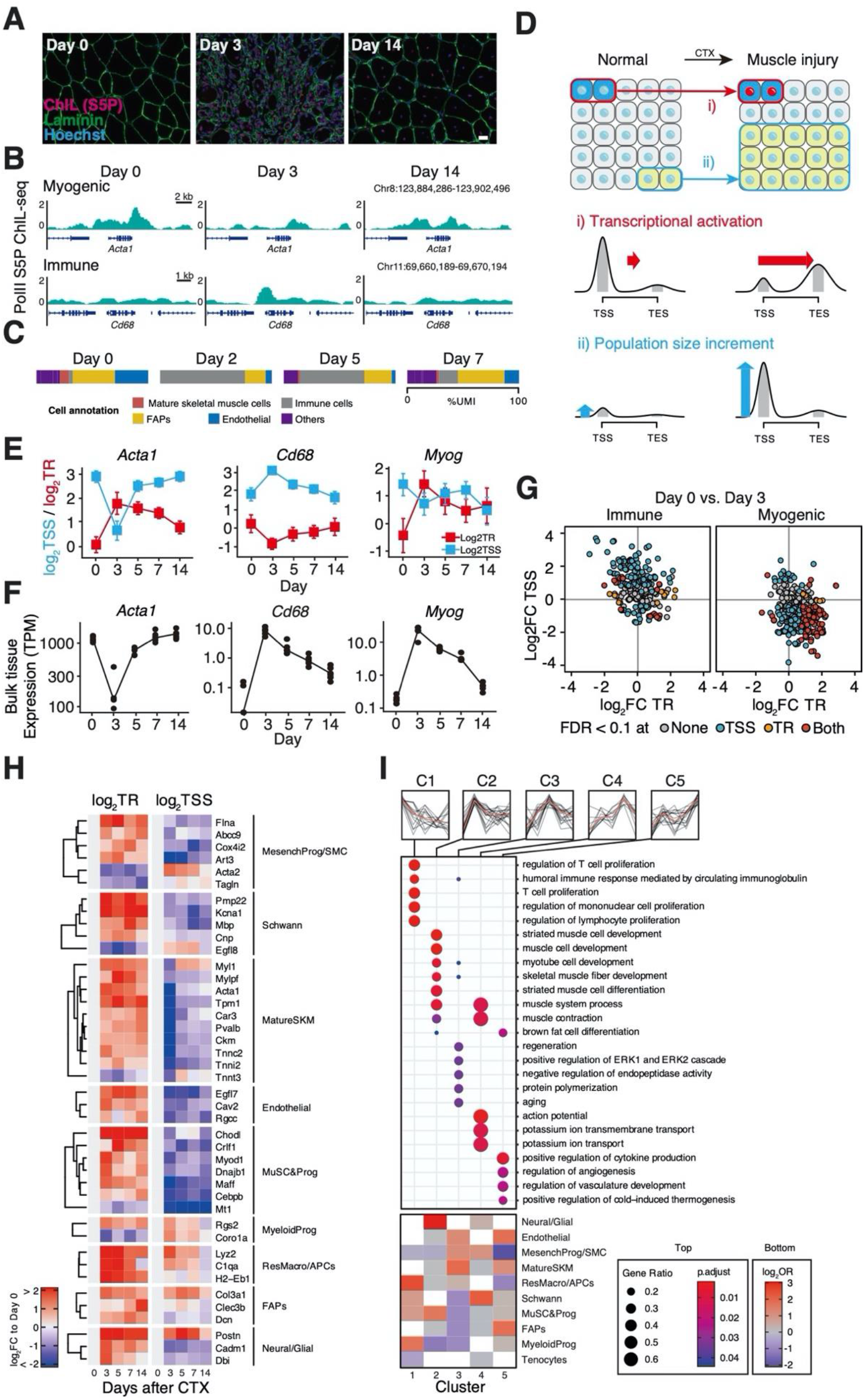
Statistical modeling of the traveling ratio reveals the independent dynamics between population and transcriptional regulation in regenerating skeletal muscle tissues. **(A)** Immunofluorescent images of the mouse tibialis anterior muscle on the indicated days after CTX treatment. The images of anti-mouse ChIL probe for RNAPII-S5P (red) and anti-rabbit IgG for laminin (green) are shown. Scale bar: 20 μm. Refer to **Figure S6A** for more frequent time points. **(B)** tsChIL-RNAPII signal of the marker genes of mature skeletal muscle (*Acta1*) and macrophages (*Cd68*). **(C)** Proportion of sequenced reads (%UMI) occupied by the representative cell types in muscle regeneration. The single cell data (GSE143437) by De Micheli et al.^48^ was re-analyzed. See **Figure S6B** for the detailed cell-type annotations. **(D)** Extraction of independent dynamics of the population and transcriptional regulation. Change in RNAPII distribution at the gene loci: a gene (blue) was transcriptionally activated (red nuclei) following the stimuli, while population size was unchanged. Change in the height of RNAPII distribution: a type of cells (yellow) was grown after the stimuli, while the transcriptional activity was maintained. **(E)** Estimated mean (95% confidence interval) of TR and the CPM of tsChIL-RNAPII at TSS. Representative genes of mature skeletal muscle cells and immune cells are shown. **(F)** Bulk-tissue expression levels (TPM) of the representative genes. **(G)** Different activities of two major cell-types in muscle regeneration. Scatter plots of log_2_FC of day-3 vs. day-0 of TR (x-axis) and the TSS-level (y-axis) are shown: immune cell marker genes (left); myogenic genes: right. Colors indicate significance in TR and TSS-level based on |log_2_FC| > 1 (two-fold) and FDR < 0.1. **(H)** Activities of major cell types in muscle regeneration. The colors of the heatmap show the log_2_FC to day 0 (uninjured) of TR and TSS levels. Representative genes with significant changes in TR are shown. **(I)** The dynamics of the biological process in muscle regeneration and the participating cell types. Genes were assigned to five groups (C1-5) based on highest time point of TR. OR indicates the specificity of participation to the biological processes.

First, we visualized the distribution of the RNAPII signal by IGV for representative genes in skeletal muscle and immune cells. Changes in RNAPII distribution are observed at the locus for *Acta1* (which is highly expressed in skeletal muscle) and *Cd68* (a surface marker of macrophages) (**Fig. 4B**). The *Cd68* locus showed an overall increase in the RNAPII signal from day 0 to day 3, whereas *Acta1* showed an overall decrease. These results indicate the rapid increase in immune cells and the decrease in skeletal muscle cells during the early stages of injury (days 2–3) as shown in **Figure 4C**. In *Acta1*, however, the RNAPII signal is more concentrated near the transcriptional end site (TES) than the transcriptional start site (TSS). We thus hypothesized that the shape of the RNAPII distribution contains information on both the population size of cells and the regulatory state of a gene known as the pause/release of the RNAPII^45,46^. Therefore, we established a model for two cases (or their combination) as shown in **Figure 4D**: one in which a specific gene of resident cells is activated by the induction of muscle regeneration (i), and the other in which the height of the already activated RNAPII signal increases due to an increase in the number of cells (e.g., migrated immune cells from outside the tissue) (ii). The traveling ratio (TR) is often used to evaluate the degree of RNAPII pause/release, as in Bartman et al.^47^, providing a brief description of the geometry of the distribution of the RNAPII in the gene loci in terms of the ratio of the signal levels between TSS and TES. Furthermore, we modeled the estimation of TR as a form of Poisson regression with an offset term (see details in Methods). For each locus, the signal level (count per million [CPM]) of RNAPII at the TSS is exp(*β_0_*), and that of TES is exp(*β_1_*) times the TSS level exp(*β_0_*), i.e., exp*(β_0_+β_1_)*. The statistical model allows us to evaluate the confidence intervals for TR and perform statistical tests for changes in varying conditions and time points.

**Figure 4E** shows the estimated values of the mean RNAPII levels at TSS and TR, along with the confidence intervals. We then compared the tissue-wide expression levels of the corresponding genes **(Fig. 4F)**. Surprisingly, the tissues-wide expression of *Acta1* and *Cd68* were synchronized with the pattern of the RNAPII TSS-level, whereas the transcription factor myogenin (*Myog*) expressed in muscle progenitor cells at the differentiation stage has a synchronized pattern to TR. These results suggest that the tissue bulk RNA-seq is a combination of the cell number and the changes in the amount of gene expression.

Therefore, to distinguish the transcriptional activation indicated by the TR, and the population size indicated by the TSS-level as inferred in **Figure 4E-F**, we analyzed the changes in the TSS-levels and TR at day 3 (**Fig. 4G** and **Fig. S6C**). Each set of genes was associated with each ‘single’ cell-type, the definition of which is based on the scRNA-seq analysis of injured muscle by De Micheli et al.^48^. The population size of the cells that express the skeletal muscle related genes (**Fig. 4G**, right) were decreased after injury, whereas the changes in TR revealed the active transcription of the genes. Meanwhile, in the group of genes associated with immune cells, TSS-level was increased while TR was less altered (**Fig. 4G**, left), which can be interpreted as an increase in the population of cells already possessing active gene loci (i.e. migration). This interpretation is consistent with the dynamic population changes in muscle regeneration clearly revealed by recent studies using scRNA-seq^48–50^. In summary, the statistical model of tsChIL RNAPII allowed us to evaluate the transcriptional activity of genes associated with specific cell types, independent of increased population of immune cells and decreased skeletal muscle cells during muscle regeneration.

Next, we identified the uncharacterized dynamics in muscle regeneration from day 0–14 using the other cell-type markers defined by De Micheli et al.^48^. First, we selected 66 genes among the markers that changed the TR (FDR < 0.1) at any time point compared with day 0. The changes in the TR and TSS level of these genes are shown as a heatmap (**Fig. 4H)** to visualize the trends in the transcriptional activation of each gene, as well as the increase or decrease in the number of cells that harbor the activated genes. From the log_2_TSS, which indicates the cell number, we confirmed that mature skeletal muscles (SKMs) decreased once after injury (white to blue); however, most genes were activated at day 3 and returned to the original population size (white) at day 14. Many of the cell types, such as mesenchymal progenitors/SMCs, myeloid progenitors, and resident macrophages/APCs, transiently increased in number after injury but returned to their pre-injured levels on day 14, indicating association with inflammatory responses (*Ada2, Rgs2, Coro1a, Lyz2, C1qa*) ^50,51^. Meanwhile, *Myl1*, a gene that was transiently increased after injury, *Tnnc2*, and *Acta1*, showed the same TR pattern, suggesting that these genes also function in regeneration and not only in muscle fiber formation^52^.

Next, we describe the muscle regeneration process by classifying gene groups according to the pattern of TR changes over time. The clusters C1-5 were assigned according to their peaks (highest point) of TR in the time-course of regeneration, the tissue-wide dynamics were appeared in **Figure 4I**, suggesting transcriptional regulation in muscle regeneration. The C1 exhibit the highest TR at day 0, and thus indicates a down regulated biological process after the injury. The proliferation of the immune cell was repressed, and the major participants are the resident macrophages and APCs and myeloid progenitors. The C2 which has peak at day 3, districted the activation of myogenesis mainly orchestrated by MuSC, muscle progenitors and also by neural cells, which is consistent with previous reports^53^. The C3, which has peak day 5, does not show strong enrichment. The C4 contained muscle contraction, ion transport and action potential related GO terms, which suggests the regenerated muscle was formed at day 7. The C5 (day 14) showed the activation of angiogenesis in the late stage of regeneration^54^. Here, the statistical modeling that combined RNAPII-mediated transcriptional elongation and population size changes achieved by our tsChIL provides a strategy for understanding the process of muscle regeneration that is organized by diverse cell types in tissue.

## Discussion

Here, we established a high-precision method for tissue epigenomic analysis using a single, thin section samples. We focused on the tsChIL data of RNAPII and established a statistical model to identify the changes in both population size and transcriptional regulation in the various cell types. In this analysis, we utilized single-cell analysis transcriptomic data as a reference of cell-type annotation. The efficient combination of existing single-cell analysis data and bulk but high-depth tsChIL data may lead to future approaches to analyze large numbers of individuals at the whole-cell level.

We demonstrated that the transcriptional regulation of each cell type can be analyzed independently, even in situations with large-scale variations in tissue cell-type composition, as in the case of muscle regeneration. tsChIL by itself can also provide a qualitative assessment of the changes in cell population size. Although we did not identify the cell types in the tissues nor estimated their compositional ratios, our framework that combined scRNA-seq and epigenomic analysis provides solid guidance for future tissue analysis.

The traveling ratio (or pausing index), a concise measure of RNA polymerase II dynamics, which was originally introduced in the ChIP-chip as a measure of the degree of transcriptional elongation^45,46^; and used in GRO-seq^55^ and ChIP-seq^56^. We found that the shape of the distribution of RNAPII at the genomic locus, as revealed by epigenomic analysis, is indeed a useful indicator of the transcriptional activity of a gene, and that the RNA-seq of bulk tissue is the sum of all transcripts of all cells and is always affected by the population size.

The statistical modeling of TR provides analogous advantages in the analysis of differentially expressed genes, such as the screening of genes with altered transcriptional states and calculation of confidence intervals for TR. Here, we used a simplified model in which the RNAPII signal at a single locus is the product of the size of the active population and the degree of activity (traveling ratio). Alternatively, a more realistic model with different transcriptional activities for different cell types and within the same cell type may be possible as proposed in the bulk data decomposition methods^22,57,58^. Despite our simplified assumption, our established model successfully determined transcriptional activities by cell type within a tissue. In addition, tsChIL RNAPII data can be modeled using a simple Poisson distribution rather than a negative binomial distribution, which involves a complex dispersion parameter estimation. Furthermore, the use of CPM normalization with offset terms as a natural way of handling replicates made the model easier to apply, interpret, and use for tissue epigenome profiling.

Conventional ChIP-seq has a limited genome coverage of cell owing to the efficiency of immunoprecipitation. In contrast, ChIL-seq, on which tsChIL is based, achieves a higher genome coverage of at least 90% for histone modifications at the single-cell level. Accordingly, the acquired data was assumed to be a sum of the deeply profiled cells. Thus, we believe that the acquisition of such high-depth epigenome data will continue to be necessary for the modeling compositions of tissues as shown in our framework. These high-depth data are expected to be provided not only by ChIL-seq, but also by other single-cell epigenomic analysis methods; thus, other methods can be integrated to our analysis framework.

tsChIL showed great potential to replace ChIP-seq, which has been the standard method of epigenomic analysis for tissues. In this paper, the high reproducibility of tsChIL, both technically and biologically, was demonstrated. Furthermore, tsChIL achieved comparable performance while using fewer cells than ChIP-seq (~1/10,000 of required cell), and parameters, such as fixation conditions, can be monitored based on the quality of immunostaining images. These advantages can reduce cost. In addition, by combining visualization and genome-wide analysis the spatial characteristic can be profiled and linked with the genome-wide characteristics of epigenomes as shown in the massive wave of RNAPII in the testis. For more advanced applications, by leveraging the pairing of serial-thin sections of the same mouse, the correlation between spatial and genome-wide patterns of heterologous proteins, such as histone modifications and transcription factors, may be reliably estimated. We believe our proposed method is a useful tool for tissue epigenomic analysis, together with recent scRNA-seq and microscopy-based spatial transcriptomics.

## Materials and Methods

### Ethical statement

All animal procedures were conducted in accordance with the Guidelines for the Care and Use of Laboratory Animals and were approved by the Institutional Animal Care and Use Committee (IACUC) at Kyushu University.

### Tissue preparation

Eight-week-old C57BL/6N mice were used as replicates for this study. The liver, left ventricle and testis were prepared from male, and tibialis anterior (TA) muscles were from female mise. Tissues were freshly frozen using isopentane chilled with LN2 and stored at −80°C. Muscle regeneration studies were performed as previously reported, except for the injection of CTX into the TA muscle^59^. Injured and intact TA muscles were sampled from five mice at day 0, 3, 5, 7, and 14 after CTX injury. The day 0 indicates a needle-injured control.

### Immunohistochemistry

Each tissue cryosection (10 μm) was placed on the bottom of 96-well microplate (Ibidi #89626) and stored at −80°C until use. Each section was fixed with 4% paraformaldehyde in 0.3% TritonX-100/PBS for 5 min and washed with 0.3% Triton X-100/PBS. Double blocking was performed using blocking one (Nacalai #03953) and M.O.M blocking reagent (Vector Laboratories #BMK-2202) following the manufacturer’s protocol. The sections were incubated overnight at 4 °C with primary antibodies diluted in M.O.M. protein concentrate/PBS, followed by incubation with ChIL probe at the same conditions but with the addition of 0.5 M NaCl. Then, the wells were filled with PBS for imaging. The following antibodies were used: rabbit anti-H3K27ac (1:500) (CMA309/9E2H10)^60^, rat anti-RNA polymerase II S5P (1:1000) (1H4B6)^61^, and rabbit anti-HNf4α (1:500) (C11F12, Cell Signaling Technology Cat. #3113), rabbit anti-laminin2α (Sigma #L-9393).

### tsChIL-seq

tsChIL-seq was performed according to ChIL^7,8^ with some modifications: longer incubation time was employed for some steps (1 h extended Tn5 binding and 2 h fill-in step), Thermo T7 RNA polymerase (100 U/well; Toyobo), and 15 cycles of polymerase chain reaction (PCR) amplification. Column purification (Zymo #D4013) and ×0.5 volume of AMpure beads (Beckman Coulter) selection were performed to obtain 200 to 500 bp average of the library. The single-end libraries were sequenced using NovaSeq (Illumina). Reads were mapped against the GRCm38 reference genome using Bowtie2^62^ with the default option. Duplicated reads were discarded using Samtools (rmdup). The uniquely mapped reads were used for further analysis.

### Quality assessments of tsChIL-seq data

The matrix of read counts on the equally sized (10 kb) windows on the mouse genome was generated using deepTools^63^ (version. 3.4.1) with the command: *multiBamSummary bins -bs 10000 --ignoreDuplicates*. Pearson correlation coefficients were calculated using the log-transformed read count (with +0.5 pseudo-counts). The breakdown of mapped reads at the genomic regions was calculated using HOMER (annotatePeaks.pl). The library complexity was evaluated by Preseq^28^. The theoretical case assumed uniform probabilities of obtaining reads from the mouse genome (i.e., a common expected values of the Poisson distribution).

### RNA-seq analysis

Total RNA (10 ng) was extracted for library preparation using a SMART-Seq Stranded Kit (Takara) according to the manufacturer’s instructions. Libraries were sequenced on Hiseq1500 and NovaSeq (Illumina). Gene expression quantification was performed using Salmon^64^ *quant* with the default option.

### Tissue-specific enhancer analysis

Peaks of tsChIL-H3K27ac were called using MACS2^65^ with the option: *callpeak --call-summits --nomodel --nolambda -q 0.05*. Tissue specificities of the peaks were evaluated using the odds ratio in the known tissue-specific enhancer lists^3^. The odds ratio is defined as *(p/(1-p))/(q/(1-q))*, where *p* is the proportion of hits in the target tissue and *q* is the proportion of hits to the other tissues in the enhancer lists. ChromVAR^66^ analysis was performed using consensus peaks of each tissue. The consensus peaks were constructed by taking the intersection of the peaks of three biological replicates. Typical and super enhancer candidates were called using HOMER^67^ *finePeaks* with the option: *-style super -superSlope -1000 -gsize 3e9*. The pre-ranked GSEA^40^ was performed using tag (read) count-ordered enhancer peaks. Then, the peaks were marked by a binary indicator overlapping with tsChIL-Hnf4a peaks (called by MACS2 as described above with the option: *-q 1e-5*).

### Transcriptional activation analysis by tsChIL-RNAPII

Aggregation plots of the gene expression percentile groups were created using agplus^68^. The gene groups were divided according to the TPM of the bulk RNA-seq analysis of each tissue (liver, heart, and testis).

### Statistical modeling of traveling ratio

The read counts of RNAPII tsChIL-seq at the TSS-region (−750 to +750 bp) and TES-region (0 to +1,500 bp) at all mouse transcripts were fitted to the following Poisson regression model. For each gene, assume that the read count *y_ij_* of the *i*-th replicate at site *j* (TSS or TES) follows the Poisson distribution, where the mean parameter *λ_ij_* satisfies the relation: *λ_ij_/M_i_: = exp(β_0_+β_1_S_ij_)*. The offset term *M_i_* is the total reads (in millions) of the replicate *i*, and *s_ij_* is the indicator variable that the read count *y_ij_* is either TSS (*s_ij_* = 0) or TES (*s_ij_* = 1). Since the offsetting is equivalent to the CPM normalization of the mean count, exp*(β_0_)* and exp*(β_1_)* can be referred to as the mean CPM at TSS and the magnification factor of TES to TSS (i.e., the traveling ratio) of the gene, respectively. The model evaluates variance and can thus estimate the confidence intervals of the traveling ratio by utilizing all replicates (5 in our case) that have different total sequenced reads. We assumed that the contrasts *X - Y* (e.g., fold-changes of TR between day 3 and day 0) follow a Gaussian distribution, and the variance was calculated from *V_X_ + V_Y_* (variances of *X* and *Y*) under the independence assumption of *X* and *Y*. *p*-values were estimated from the model, and multiple test correction was performed using the Benjamini-Hochberg procedure in the selected genes of interest.

## Data Availability

The RNA-seq and tsChIL-seq data generated in this study have been deposited in the Gene Expression Omnibus (GEO) database under the accession code: GSE159024. The codes used for the statistical modeling of tsChIL-seq data are available at: https://github.com/kazumits/tissueChIL

## Author Contributions

K.M., K.To., A.H. and Y.O. conceived and designed the experiments. K.Ta. and A.H. performed the experiments. K.M. and K.Ta. analyzed the data. K.M. performed statistical analysis. S.S., S.O., T.H., H.Ku. and H.Ki. contributed materials and analysis tools. K.M., K.To. and Y.O. wrote the paper. All authors read and approved the final manuscript.

## Acknowledgments

Computations were carried out using the computer resources offered under the category of Intensively Promoted Projects by the Research Institute for Information Technology at Kyushu University. We appreciate the technical assistance from The Research Support Center, Research Center for Human Disease Modeling, Kyushu University Graduate School of Medical Sciences. This work was in part supported by JST PRESTO JPMJPR2026 to K.M., JPMJPR19K7 to A.H., JST CREST JPMJCR16G1 to Y.O., H.Ku. and H.Ki.; JST ERATO JPMJER1901 to H.Ku.; MEXT/JSPS KAKENHI JP19H04970, JP19H03158 and JP20H05393 to K.M.; JP18K19432, JP19H03211, JP19H05425 and JP20H05368 to A.H.; JP18H05534 and JP20H00449 to H.Ku.; JP18H04802, JP18H05527, JP19H05244, JP17H03608, JP20H00456 and JP20H04846 to Y.O.; JP18H05527 and JP17H01417 to H.Ki.; AMED JP20ek0109489h0001 to Y.O., AMED BINDS JP19am0101076 and JP20am0101076 to H.Ku.; JP19am0101105 to H. Ki.

## Supplementary Information

**Figure S1:**
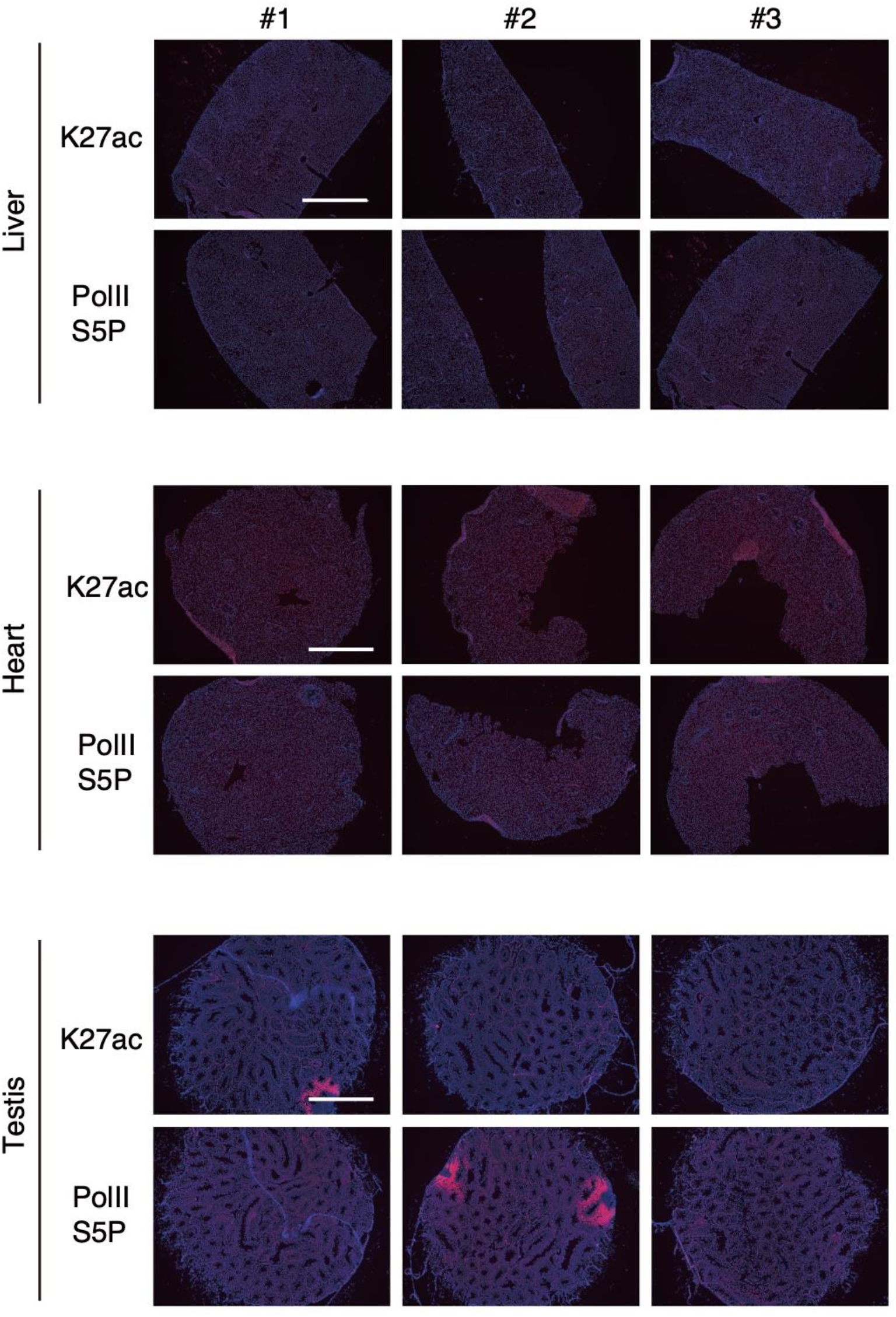
Immunofluorescent images of whole sections stained with the ChIL-probe. Immunofluorescent images of the indicated tissues for all replicates (*N*=3). Tissue sections were stained with H3K27ac or PolIIS5P antibody and visualized using the fluorescent dye–conjugated ChIL-probe. DNA was counterstained with Hoechst 33342. Scale bar: 1 mm.

**Figure S2:**
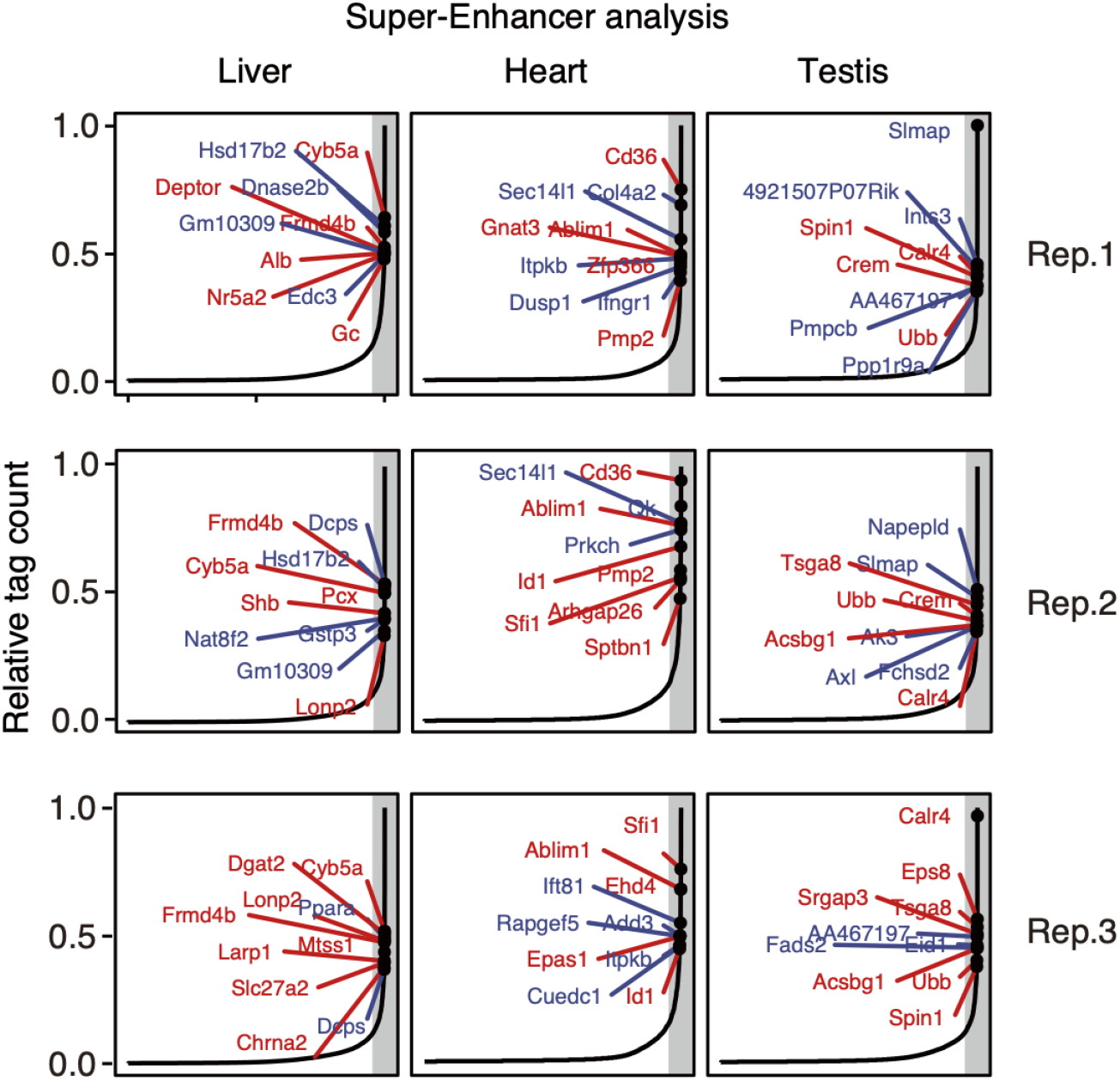
Super-enhancer analysis of each replicate. Tissue-specific enhancers are identified so that they are listed more than twice (twice: blue, all: red) in the top 5% of tag count among enhancer candidates and are not in the SEs of other tissues. Grey shades indicate the top 5% of tag count among the enhancer candidates.

**Figure S3:**
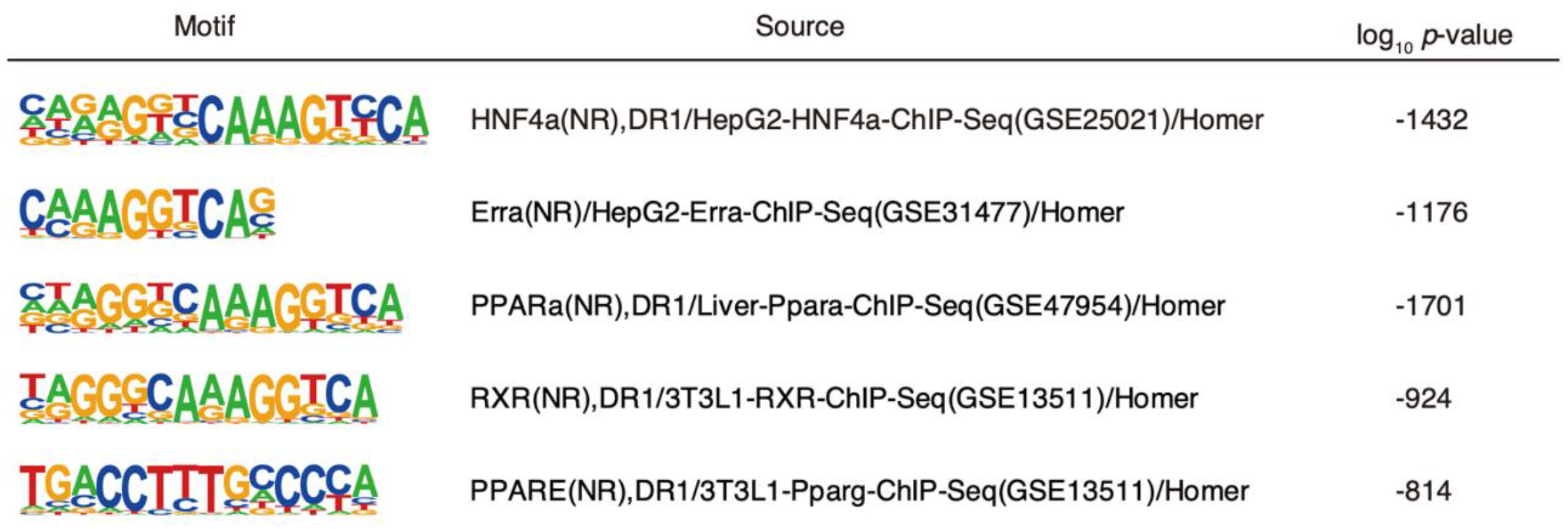
Motif enrichment analysis of tsChIL-Hnf4a peaks. Enrichment analysis of *known motifs* using HOMER. The motifs shown here are the top 5 based on the *p*-values. Their enrichment of motifs was evaluated within 250 bp from a summit of MACS2 peaks. The height of the motif logos corresponds to nucleotide frequencies.

**Figure S4:**
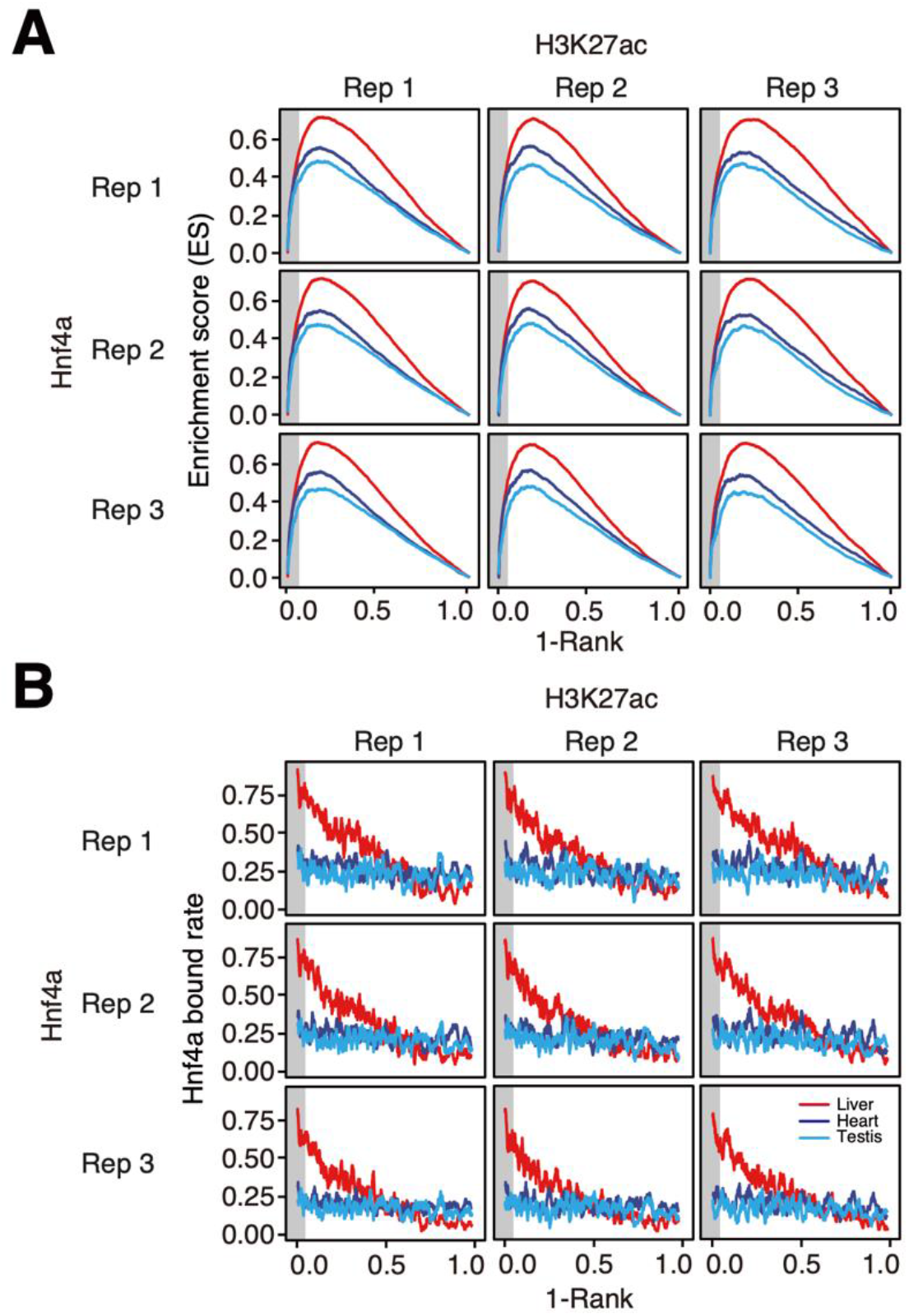
Enhancer set enrichment analysis of Hnf4a-bound genes. Gene set enrichment analysis of Hnf4α-bound genes **(A)**, and the rate of Hnf4α-bound genes in the sliding windows of 100 genes **(B)** along the ordered enhancers. All possible combinations (3 × 3 combination of replicates for tsChIL-H3K27ac and tsChIL-Hnf4α) are shown.

**Figure S5:**
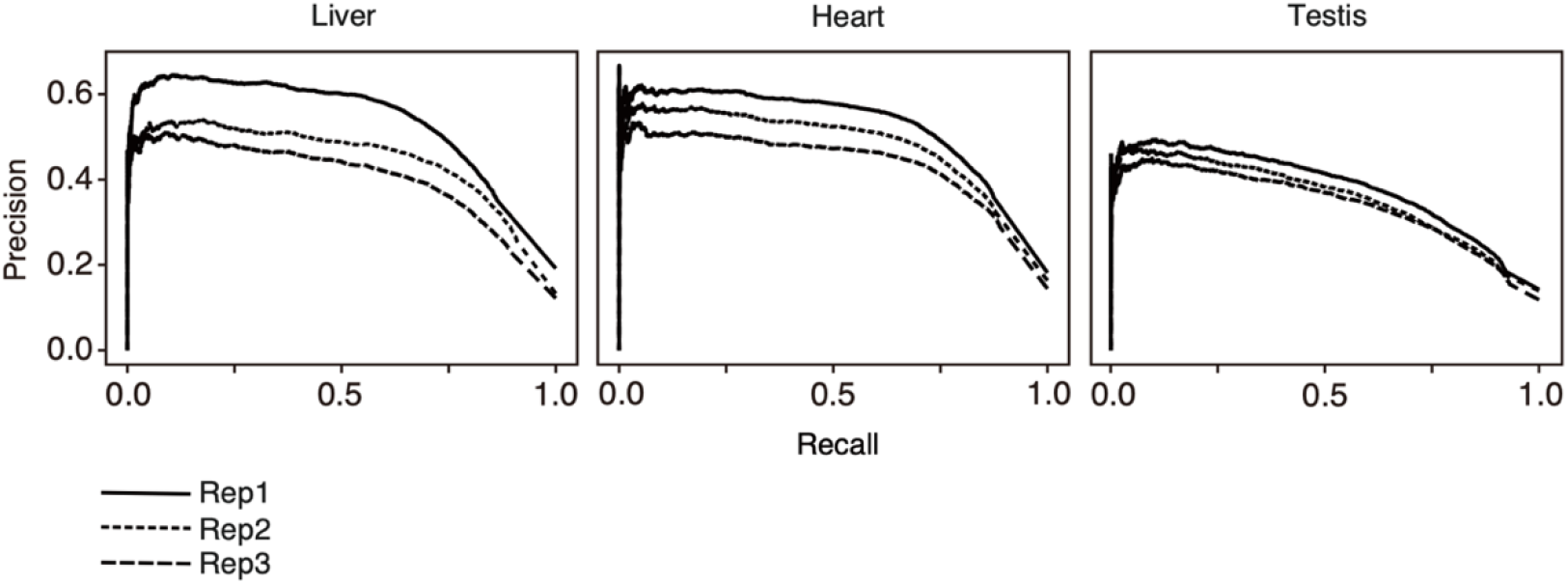
Tolerant definition of “active genes” by RNA-seq. Precision and recall curves for predicting tsChIL-RNAPII peaks based on TPM values are shown. The recall represents the proportion of RNAPII peaks covered by the active genes, and the precision is the proportion of active genes covered by the RNAPII peaks. Active genes are defined at each TPM threshold.

**Figure S6:**
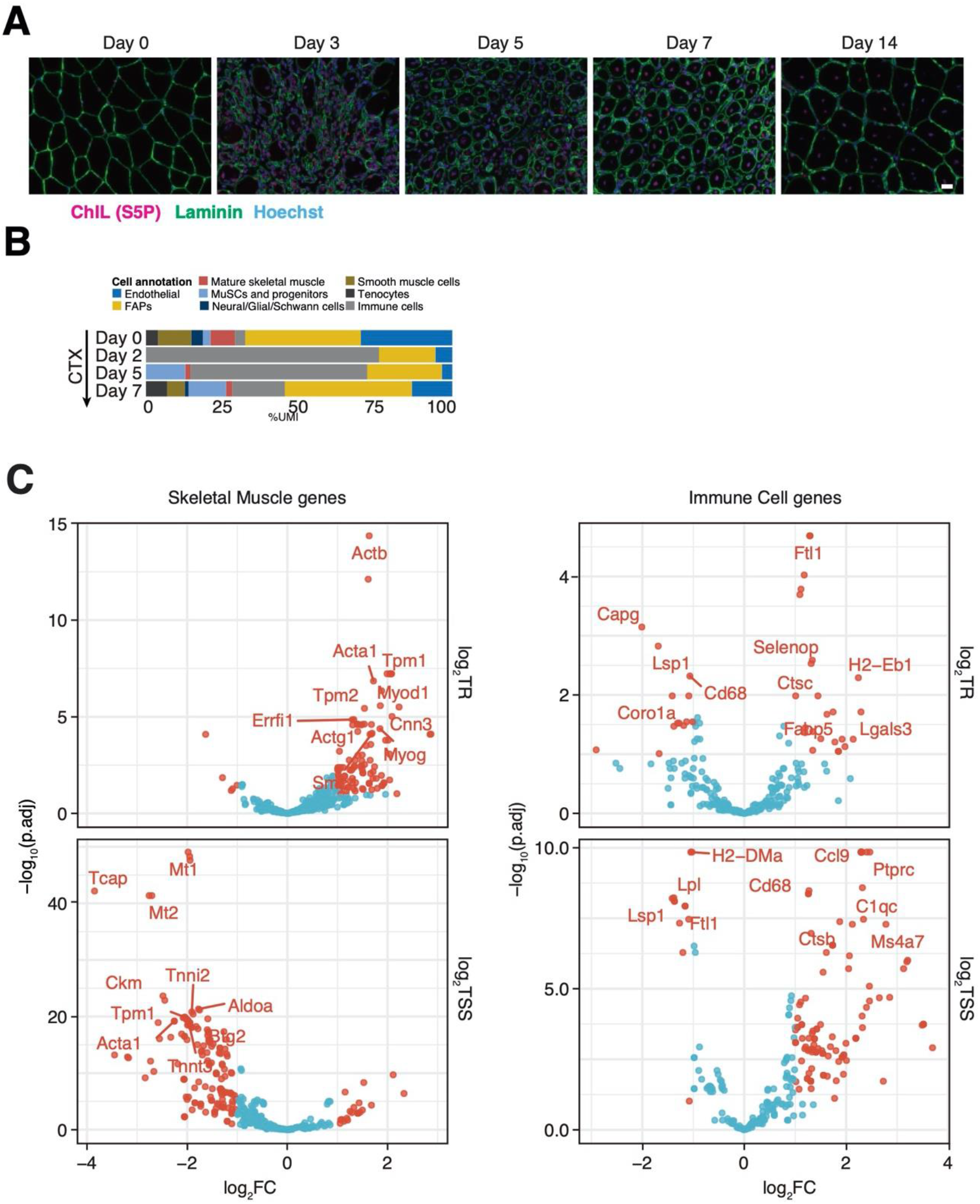
Statistical modeling of RNAPII reveals transcriptional dynamics in muscle regeneration. **(A)** The complete immunofluorescent images shown in Figure 4a. Scale bar: 20 μm. **(B)** Proportion of sequenced reads (%UMI) occupied by the annotated cell types in muscle regeneration. The single cell data (GSE143437) by De Micheli et al. was re-analyzed. **(C)** Volcano plots of the contrasts (day 3 vs. day 0 after CTX injury) for TR (top) and TSS (bottom). The x-axis represents log_2_FC (day 3/day 0), whereas the y-axis represents −log_10_FDR. Significant changes that satisfy |log_2_FC| > 1 (twofold) and FDR < 0.1 are in red. Genes that have the top 10 *p*-values are labelled.

**Table S1:**
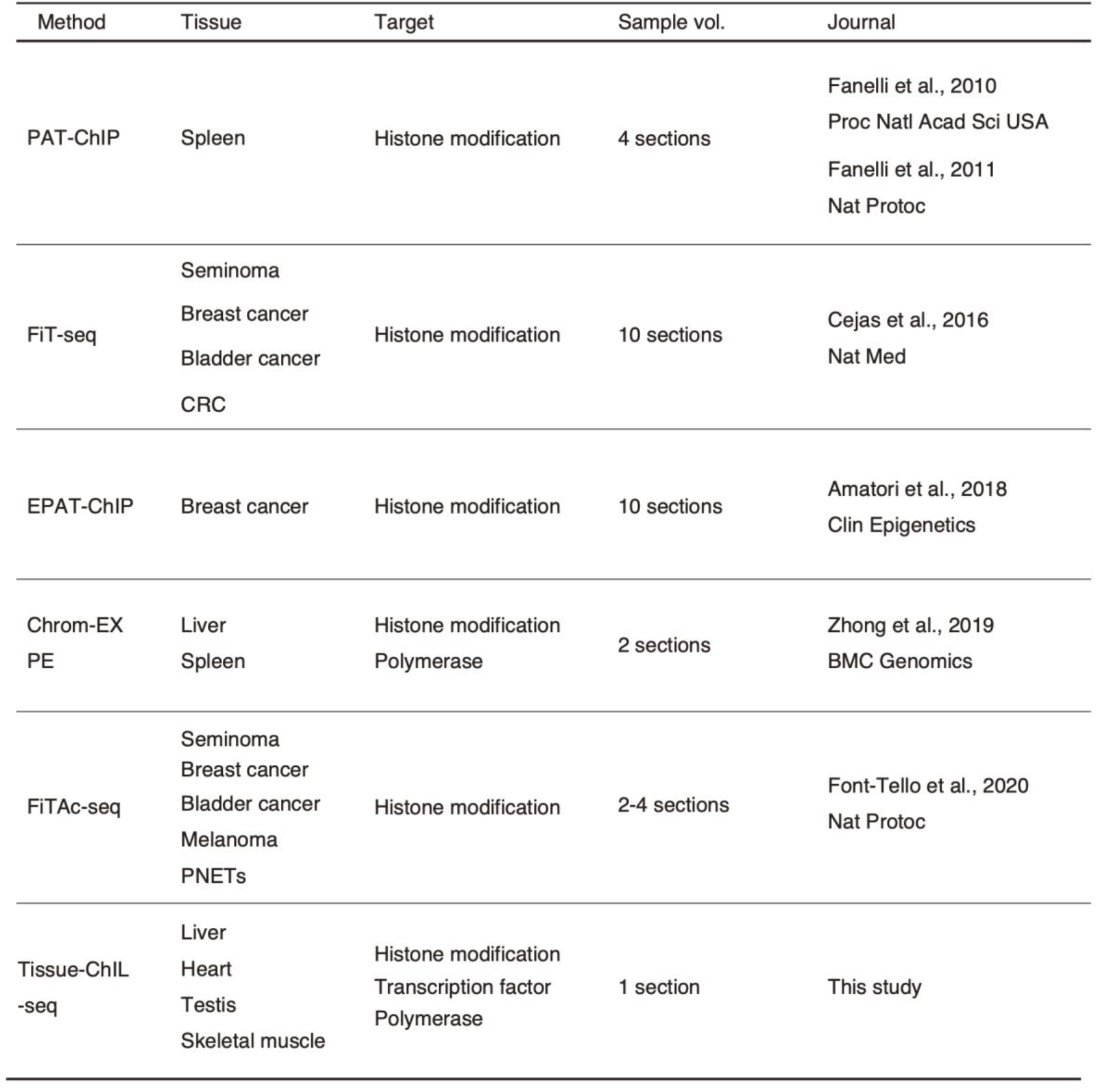
Epigenomic analysis methods on tissue sections.

